# Desiccation stress enhances virulence potential of group B *Streptococcus*

**DOI:** 10.64898/2026.11.10.765798

**Authors:** Tamanna Jafar, Jéssica da Conceiçao Mendonça, Nicole Boland, Lindsey R. Burcham

## Abstract

*Streptococcus agalactiae* (Group B *Streptococcus*, GBS) asymptomatically inhabits the gastrointestinal and female reproductive tracts but can also cause invasive diseases in susceptible individuals. Despite its clinical importance, relatively little is known about its ability to survive outside the host and whether environmental persistence contributes to transmission and infection. Here we investigate the GBS survival under nutrient starvation and desiccation, characterizing phenotypic and genomic adaptations associated with environmental persistence. GBS remained viable for 30 days during starvation at room temperature and for more than 80 days under desiccated conditions, with GBS grown to stationary-phase bacteria exhibiting greater survival than actively growing bacteria. Survival under desiccation was conserved among several clinically relevant GBS serotypes. Following rehydration desiccated GBS exhibited enhanced biofilm formation compared to control non-desiccated strains. Whole-genome sequencing identified mutations in genes involved in capsule biosynthesis, cell-envelope homeostasis, nutrient transport, and carbohydrate metabolism, suggesting that multiple adaptive pathways may contribute to environmental persistence. Importantly, desiccated GBS retained the capacity to colonize the vaginal mucosa and displayed increased invasion of reproductive tissues in a murine vaginal colonization model. In a systemic infection model, desiccated GBS exhibited enhanced virulence compared to control non-desiccated GBS, demonstrating that environmental survival does not diminish pathogenic potential. This environmental persistence may represent an underappreciated factor contributing to GBS transmission and the epidemiology of late-onset neonatal disease.

**Importance:** Group B *Streptococcus* is a leading cause of neonatal sepsis and meningitis, particularly during late-onset disease for which the routes of transmission remain poorly understood. Although GBS is generally considered a host-associated pathogen, increasing epidemiological evidence suggests that environmental sources may contribute to transmission. However, little is known about the ability of GBS to survive under environment conditions different than those encountered within a host. In this study, we demonstrate that GBS withstands desiccation and nutrient deprivation for long periods while retaining its infectivity *in vivo*. These findings identify environmental persistence as a previously unexplored feature of GBS and suggest environmental reservoirs may contribute to GBS transmission, particularly in health care settings. Understanding the mechanisms that enable GBS to persist outside the host may improve current strategies to prevent infections and help in reducing the burden of GBS disease.

## Introduction

*Streptococcus agalactiae*, or Group B *Streptococcus* (GBS), is a Gram-positive bacterium that colonizes up to 30% of healthy women within the gastrointestinal and genitourinary tracts [1-3]. GBS vaginal colonization can result in complications during pregnancy such as preterm labor, preterm premature rupture of membranes, chorioamnionitis, and stillbirth [3, 4]. Maternal rectovaginal colonization of GBS is the primary reservoir for neonatal GBS transmission during labor and delivery, leading to neonatal sepsis, pneumonia, and meningitis, which are associated with substantial morbidity and mortality [5, 6]. Globally, an estimated 19.7 million pregnant women were colonized with GBS in 2020, resulting approximately 58,300 infant deaths due to invasive GBS disease [7].

Neonatal GBS disease is broadly classified into early-onset disease (EOD) and late-onset disease (LOD) [8]. EOD, which occurs within the first six days of life, is primarily acquired through vertical transmission during delivery. Whereas LOD, occurring between 7 days and 3 months of age, is thought to result from postnatal exposure to GBS [3]. Globally, approximately 231,800 EOD cases and 162,200 LOD cases were estimated to have occurred in 2020, corresponding to 59% and 41%, respectively invasive infant GBS cases [7]. Although the incidence of EOD has been significantly reduced through the implementation of intrapartum antibiotic prophylaxis (IAP), IAP has had little impact on cases of LOD, which remains a significant cause of neonatal morbidity and mortality [2, 6]. Consequently, in settings with widespread IAP implementation, the relative burden of LOD has become increasingly prominent [9, 10]. In the United States, a 2024 surveillance estimated EOD and LOD incidences of 0.15 and 0.2 cases per 1000 live births, respectively [11]. In most cases, the reservoirs of LOD remain unidentified but, potential sources of LOD include maternal skin, breastmilk, exposure to contaminated environmental surfaces, and poor hygiene practices [12]. LOD is also associated with prolonged hospitalization, transmission between infant siblings from multiples, frequent invasive procedures, and extended admission to neonatal intensive care units (NICUs), particularly among premature and low birthweight infants who are highly susceptible to infection [13-16].

While advances in neonatal intensive care have improved the survival of preterm infants, they have also increased the risk of nosocomial infections, with GBS remaining one of the leading causes of neonatal sepsis [12, 13, 17]. Recent epidemiological studies have reported that hospital-associated clusters of LOD have been reported with prolonged intervals between cases, supporting the possibility that environmental persistence and horizontal transmission may contribute to neonatal infection in clinical settings [18-20]. However, despite increasing reports supporting healthcare-associated and environmental transmission, the ability of GBS to survive under prolonged abiotic conditions has remained largely unexplored. Therefore, we aim to investigate how GBS survives extended nutrient limitation and environmental desiccation and examine how stress adaptation impacts GBS virulence potential.

## Results

### GBS survives prolonged nutrient deprivation and desiccation stress

To investigate the ability of GBS to withstand prolonged nutrient limitation and determine how temperature influences persistence, GBS grown to logarithmic phase (OD_600nm_ 0.4) and stationary phase (OD_600nm_ 0.8) were incubated at 4°C, 25°C, and 37°C in PBS. At 4°C, bacterial viability remained nearly unchanged from the initial inoculum after 30 days of starvation **(Figure 1A)**. Cultures incubated at 25°C exhibited a gradual decline in viability throughout the experimental period. However, bacteria remained viable through more than 30 days of nutrient limitation **(Figure 1B)**. In contrast, GBS viability from both growth phases declined rapidly when cultures were stored at 37°C, with no viable GBS recovered after day 18 **(Figure 1C)**. For both experiments performed at 25°C and 37°C, cultures grown to stationary phase exhibited more resilience to starvation than cultures grown to log-phase.

**Figure 1:**
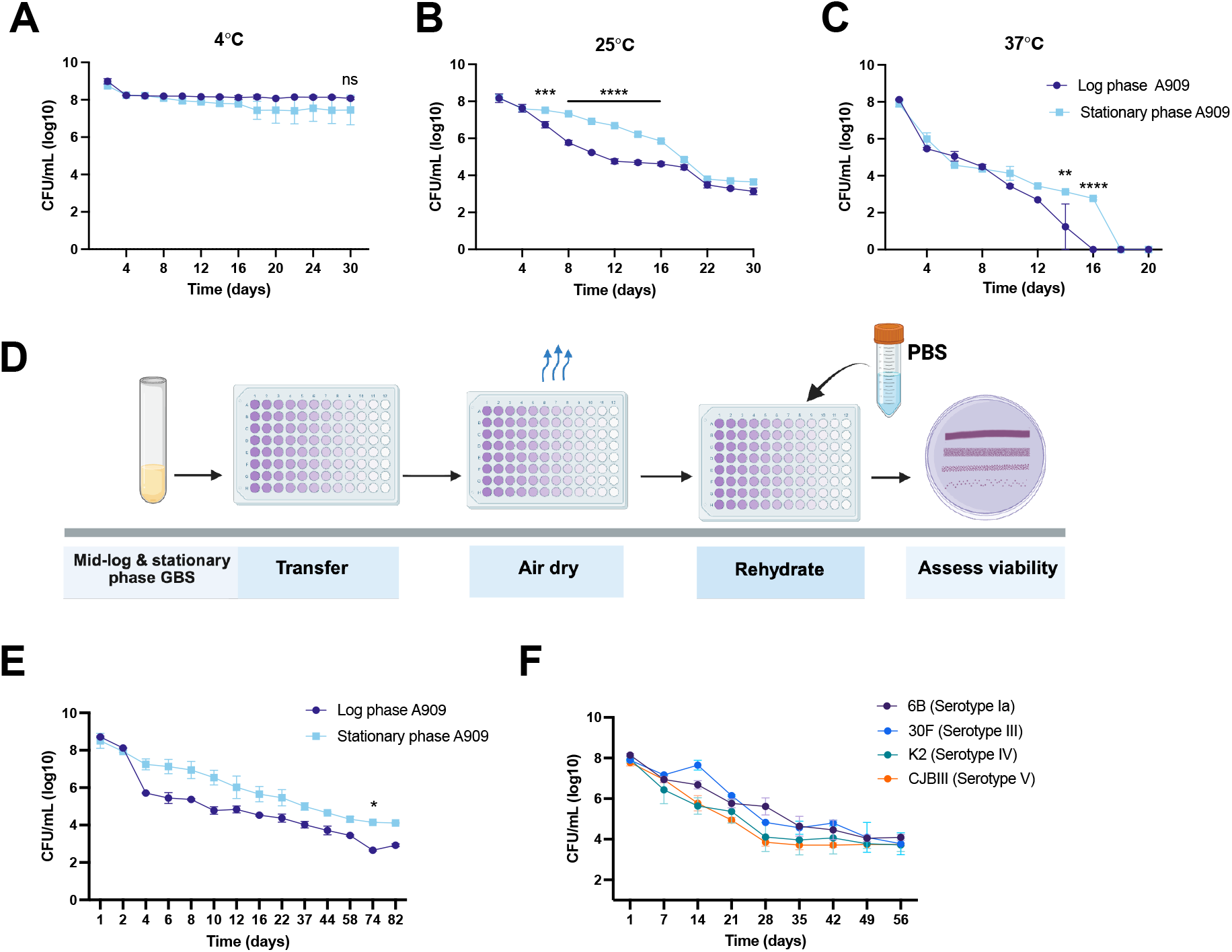
GBS survives prolonged nutrient deprivation and desiccation stress. Survival of log and stationary phase A909 during nutrient deprivation at **A)** 4°C, **B)** 25°C, and **C)** 37°C. **D)** Schematic of the desiccation protocol. Survival of desiccated **E)** A909 or **F)** GBS mid-log phase clinical vaginal (6B, 30F, K2) and systemic (CJBIII) isolates for extended time courses. Data are presented as mean ± SEM from two independent biological replicates. Statistical analyses were performed using two-way ANOVA with Sídák’s multiple-comparisons testing. Statistical significance is indicated as *p \**< 0.05, **\*\***< 0.01, ***< 0.001, and ****< 0.0001.

Given the potential role of environmental persistence in healthcare-associated transmission, we aimed to determine how effectively GBS survives low-moisture conditions that it may encounter on abiotic surfaces. GBS cultures were dried in 96-well plates, stored at room-temperature, and were rehydrated at individual timepoints throughout the experimental timeline **(Figure 1D)**. Cultures of GBS grown to logarithmic phase and stationary phase retained viability through more than 80 days of desiccation **(Figure 1E)**. Viability gradually declined over time in cultures seeded from both growth states, and a slight but statistically significant difference was observed between stationary phase and log-phase grown cultures on day 74 post-desiccation. To determine whether desiccation tolerance was reproducible across genetically diverse GBS strains, we tested survival during desiccation stress in clinical isolates of capsular serotypes Ia, III, IV, and V, and across all strains, viable bacteria were recoverable for more than 50 days, with all cultures remaining stable around a burden of 1×10^4^-1×10^5^ CFU/mL **(Figure 1F)**. These results highlight the ability of GBS to withstand prolonged desiccation stress is reproducible across a diverse range of clinical isolates regardless of capsular serotypes. Overall, these findings demonstrate that GBS is capable of long-term persistence during nutrient deprivation and desiccation.

### Desiccation stress promotes GBS biofilm formation

To determine whether desiccation influences the ability of GBS to form biofilms, we performed crystal violet staining to quantify biofilm formation produced by non-desiccated GBS A909 strain and a strain desiccated for 30 days (D30). The D30 strain produced significantly greater biofilm biomass than the non-desiccated control **(Figure 2A)**. Furthermore, biofilms formed by both strains were examined by confocal laser scanning microscopy. Images from the desiccated D30 strain showed increased biofilm density compared to the non-desiccated control A909 **(Figure 2B)**, that was quantifiable by calculating mean fluorescence intensity **(Figure 2C)**. We tested additional strain backgrounds including GBS vaginal clinical isolates representing Serotypes Ia (6B), III (30F), and IV (K2), and a serotype V invasive isolate (CJBIII), and their paired desiccated strains. Like A909, the serotype III 30F strain exhibited a trend towards increased biofilm formation following desiccation (*p=0*.*06)*, while desiccation appeared to have minimal impact or a low level of inhibition for other clinical strains, though these did not reach statistical significance **(Figure S1A)**. To determine whether desiccation altered additional virulence-associated phenotypes, β-hemolysin/cytolysin production, adherence and invasion of VK2 vaginal epithelial cells, and tolerance to environmental stressors were evaluated. Desiccation did not affect hemolytic potential **(Figure S1B)**, adherence to VK2 cells **(Figure S1C)**, or susceptibility to general cell stress induced by UV, ethanol, or sodium hypochlorite **(Figure S1D-F)**, although the GBS desiccated strain D30 exhibited reduced invasion of VK2 cells compared with the non-desiccated A909 control strain (**Figure S1G)**.

**Figure 2:**
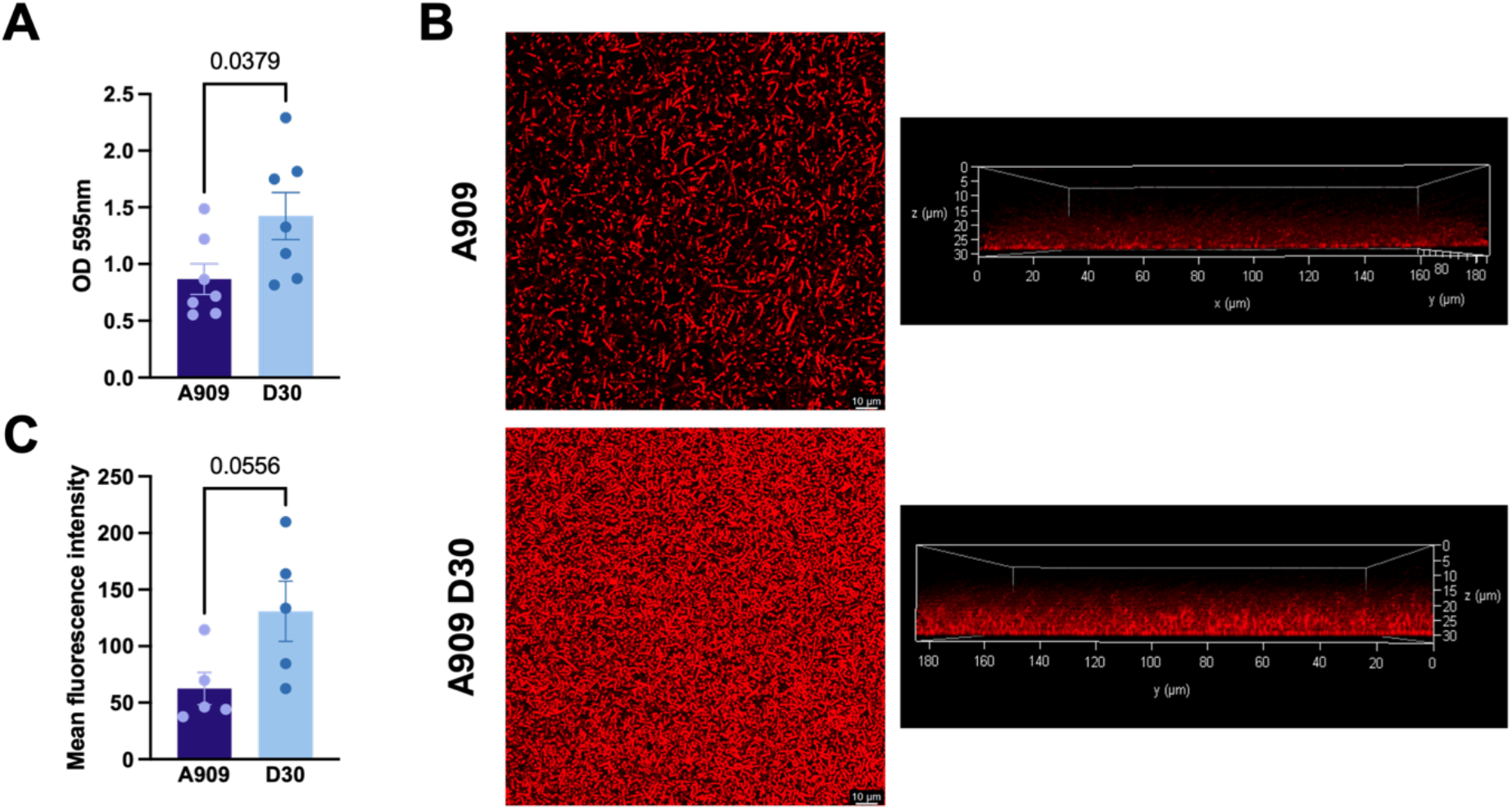
Desiccation stress promotes GBS biofilm formation. **A)** Crystal violet biofilm assay comparing wild-type non-desiccated A909 and D30 GBS. **B)** Representative confocal microscopy images. **C)** Quantification of biofilm biomass by ImageJ measuring mean fluorescence intensity from 63x confocal images. Data are presented as mean ± SEM from at least 5 independent biological replicates. Statistical analyses were performed using the nonparametric Mann–Whitney test. Calculated *p* values are indicated on the respective figures.

### Desiccation enhances GBS fitness *in vivo*

A prior study determined that desiccated *S. pneumoniae* remain infectious following rehydration and can establish in the nasopharynx, demonstrating that environmental persistence can occur without loss of the capacity to colonize a host [21]. To determine the consequences of desiccation on GBS colonization, persistence, and ascension of the reproductive tract, we used a previously established murine vaginal colonization model [22]. Female 8-week-old mice were synchronized in estrus with an injection of 0.5mg/100μL 17β-estradiol and 24 hours later intravaginally inoculated with the non-desiccated GBS strain A909 or the desiccated D30 strain **(Figure 3A)**. Vaginal swabs collected on days 1 and 4 post-colonization confirmed that both GBS strains colonized the vaginal lumen, with no significant differences in bacterial burden recovered from animals colonized with either strain **(Figure 3B)**. However, Fisher’s exact test showed that animals colonized with the non-desiccated GBS had higher odds of clearing GBS, compared to animals colonized with the desiccated GBS strain (*p*=0.0187, **Figure 3C**). On day 5 post-colonization, murine reproductive tissues were harvested, homogenized, and plated to further evaluate tissue invasion and ascending infection. We identified significantly higher bacterial burdens recovered from vaginal, cervical, and uterine tissues of animals colonized with the desiccated D30 strain compared to those colonized with the non-desiccated A909 strain (**Figure 3D-F)**. These findings suggest that following desiccation stress, GBS retains infectivity and exhibits enhanced ascension and invasion of the reproductive tract tissues.

**Figure 3:**
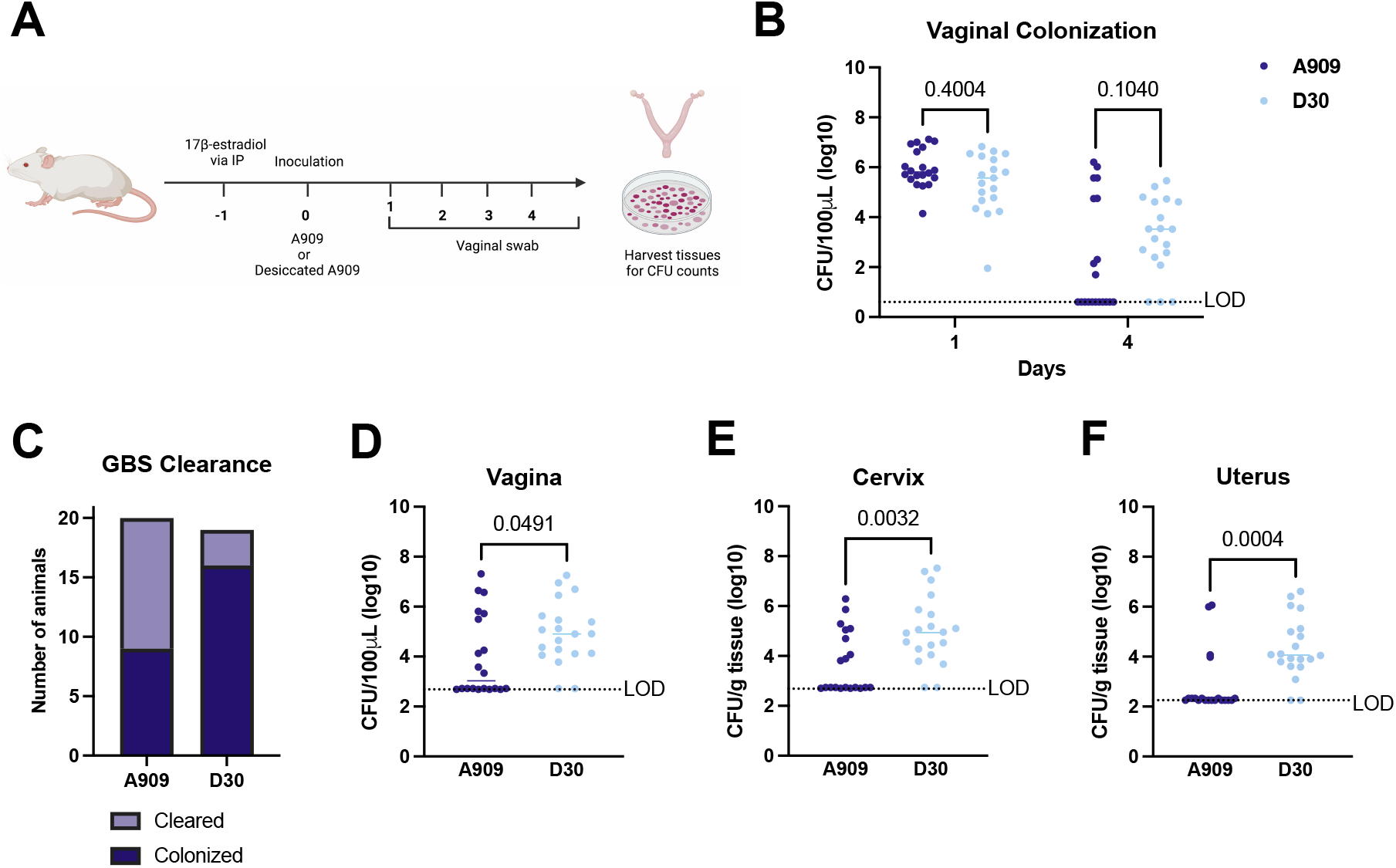
Desiccation enhances GBS ascending infection. **A)** Experimental design of the murine vaginal colonization model. **B)** Luminal colonization of mice inoculated with A909 or A909 D30. **C)** Data stratified by animals that cleared GBS infection versus those that remained colonized by either non-desiccated or desiccated GBS. Bacterial burden recovered from tissue homogenates of the **D)** vagina, **E)** cervix, **F)** and uterus of animals harvested on day 5 post-colonization. Data are pooled from the two independent experiments with each symbol representing one animal with a total of 39 animals (20 colonized by non-desiccated GBS and 19 colonized by desiccated D30 GBS) included. The horizontal bars represent the data median **(B-F)**. Statistical analyses of colonization data were calculated by two-way ANOVA with Sídák’s multiple comparisons test, comparison of clearance versus colonization were calculated by Fisher’s exact test, and tissue burdens were analyzed by Welch’s t-test. Calculated *p* values are indicated on the respective figures.

As postnatal acquisition of GBS poses a significantly threat to the neonate and is capable of progression to invasive late onset disease (LOD) including sepsis and meningitis, we next examined whether GBS surviving desiccation retained virulence in a hematogenous model of bloodstream infection. To test this, female 8-week-old mice were inoculated retro-orbitally with the non-desiccated control strain, A909 and the desiccated D30 strain. Animals were monitored for signs and symptoms of systemic infection up to 83 hours post-infection. Animals infected with the desiccated GBS strain displayed increased disease severity, but this did not ultimately result in more profound mortality throughout the course of infection **(Figure 4B-C)**. At the experimental endpoint, animals were humanely euthanized and blood and systemic tissues were harvested, homogenized, and plated for bacterial burden. Significantly higher bacterial burden was recovered from both blood and brain tissues of animals infected with the D30 desiccated strain compared to those challenged with the non-desiccated control A909 strain **(Figure 4D-E)**. Together, these findings recapitulate what was observed in the vaginal colonization and ascending infection model and suggest that GBS not only retains infectivity following desiccation, but that desiccation stress may prime GBS virulence potential or increase resistance to the immune response.

**Figure 4:**
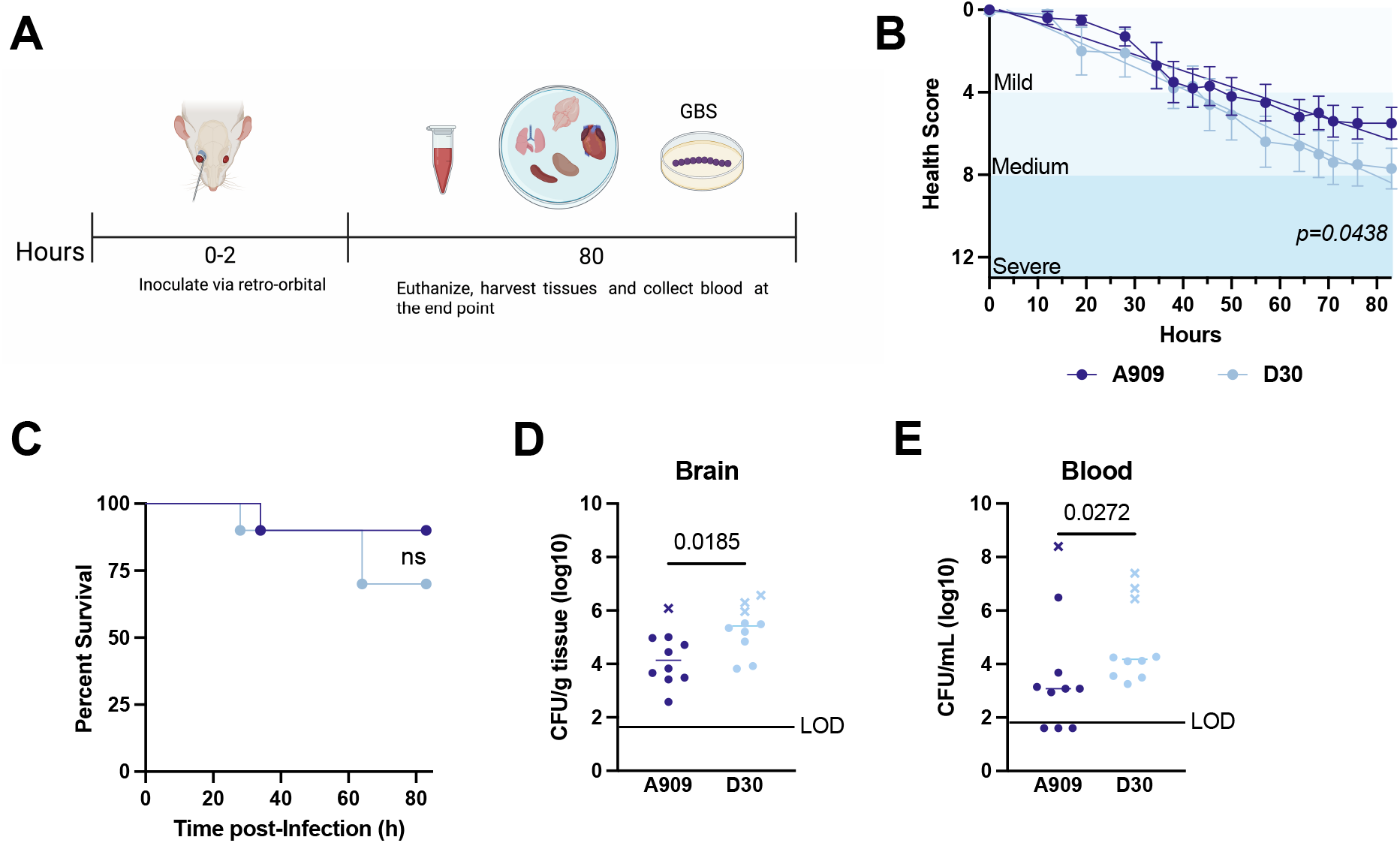
Desiccation exacerbates GBS systemic infection. **A)** Experimental design of the murine systemic infection model. **B)** Clinical disease progression following infection with non-desiccated and desiccated GBS. **C)** Percent survival of animals throughout the course of infection to the experimental endpoint. Bacterial burdens recovered from the **D)** brain and **E)** blood harvested at the time of moribund (denoted by x symbol) or the experimental endpoint of 83 hours post-infection (denoted by circle). Each symbol represents one animal with a total of 20 animals (10 colonized by non-desiccated GBS and 10 colonized by desiccated D30 GBS) included and horizontal bars mark the median of the data **(D&E)**. Statistical analysis of brain and tissue burden was performed using a Mann–Whitney test and calculated *p* values are indicated on the respective figures.

### Whole genome sequencing reveals genetic variation in desiccated strains

To identify potential genetic alterations between non-desiccated control GBS and the populations that survived desiccation stress, we performed whole-genome sequencing and comparative genomic analyses. We identified multiple sequence variants in the desiccated population, including single nucleotide polymorphisms (SNPs), deletion and insertion events, and intergenic mutations distributed throughout the genome that were absent in the non-desiccated strain. Among the coding variants, SNPs were identified in genes encoding proteins involved in nutrient transport, cell envelope homeostasis, metabolism, and surface-association. These included mutations in a putative nickel ABC transporter substrate-binding protein (NikA), a HIT family protein, PTS sugar transporter subunit IIA, cell wall-active antibiotics response protein (LiaF), Penicillin Binding protein 3 (PBP3), glycogen synthase (GlgA), DUF1149 family protein, D-alanine--D-alanine ligase (Ddl), and LPXTG cell-wall-anchored fibrinogen-binding protein (FbsA). Two single-base deletion events and a four-base insertion event were detected that resulted in frameshift mutations within the coding region of the capsular polysaccharide synthesis protein CpsH. Several mutations were also identified in intergenic regions, including loci between SAK_RS06710 and *rplL*, which encodes the 50S ribosomal protein L7/L12 involved in protein synthesis; between SAK_RS11685 and SAK_RS06750, adjacent to a TetR-family transcriptional regulator; between SAK_RS11910 and SAK_RS11460, adjacent to a hypothetical protein; and upstream of *infB*, which encodes the translation initiation factor IF-2 required for initiation of protein synthesis (**Table 1**).

**Table 1.**
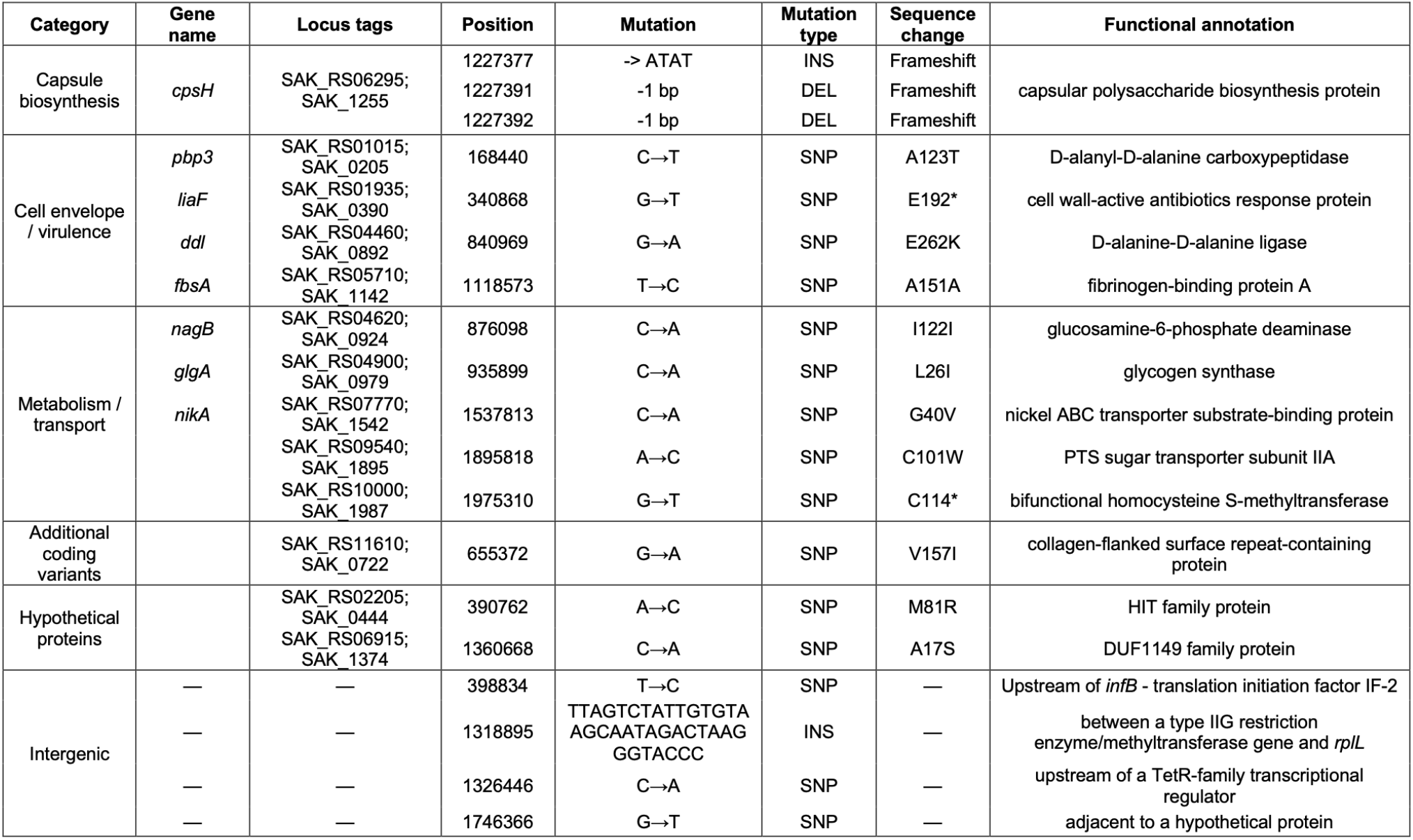
Mutations detected in D30 population compared to non-desiccated A909 control.

## Methods

### Animal ethics statement

Animal studies were performed at the University of Tennessee, an AAALAC accredited institution, in accordance with the Institutional Animal Care and Use Committee (IACUC) under protocol # 2937-1222. All experiments were performed under the approval of the Institutional Biosafety Committee protocol no. 22-580-2.

### Bacterial strains and growth conditions

*Streptococcus agalactiae* (GBS) neonatal clinical isolates A909 (serotype Ia) and CJBIII (serotype V), as well as GBS vaginal colonization isolates K2 (serotype IV), 30F (serotype III), and 6B (Serotype Ia), were used in this study. Vaginal clinical isolates were isolated from vaginal swabs collected from term-pregnant individuals at the time of admission to labor and delivery under the protocol no. 5129 approved by the University of Tennessee Health Science Center College of Medicine, Knoxville Institutional Review Board. GBS strains were grown for 16-18 hours in Todd-Hewitt Broth (THB, BD Diagnostics) at 37ºC overnight prior to subculture in fresh THB. For all strains, growth was monitored by measuring the optical density at 600 nm (OD_600nm_).

### Nutrient starvation and desiccation tolerance

GBS A909 was grown overnight in THB at 37°C, sub-cultured in fresh THB, and incubated until reaching mid-log phase (OD_600nm_ 0.4, ∼1 x 10^8^ CFU/mL). Mid-log cultures were pelleted and normalized to an OD_600nm_ of 0.4 in 1x PBS. For stationary phase cultures, overnight cultures were pelleted and normalized to an OD_600nm_ of 0.4 in 1x PBS. Mid-log and stationary normalized cultures were sealed with parafilm and stored at 4°C, 25°C, or 37°C until samples were collected, serially diluted, and plated on THB agar for CFU/mL enumeration at each time point. For experiments studying desiccation tolerance, normalized mid-log and stationary phase cultures were aliquoted at 25 μL per well into a 96-well plate, and plates were air-dried in a biosafety cabinet with airflow for 24-hours, and were stored in the biosafety cabinet at room temperature for the duration of the experiment. Individual wells were assigned to each experimental time point and were rehydrated at the time of sample collection. At each time point the designated wells were rehydrated with 25 μL PBS, bacterial cells were mixed thoroughly by pipetting and scraping, and samples were serially diluted and plated on THB agar to monitor GBS viability. Recovered bacteria were collected, outgrown in THB, and frozen in glycerol at -70°C on day 30 post-desiccation and were used in downstream analyses to represent the desiccated population. Clinical isolates were outgrown and frozen at -70°C on day 56 post-desiccation.

### Biofilm assay

Frozen stocks of non-desiccated and desiccated strains were cultured overnight and sub-cultured the morning of each experiment to mid-log growth phase. Strains were normalized as described above in PBS before inoculation. Each strain was diluted 1:100 in fresh THB medium to obtain an initial inoculum of approximately 1 x 10^6^ CFU/mL, and 200 μL was added per well in a minimum of three technical replicates. After 48 hours, the plate was washed once with PBS to remove planktonic cells. Then the adherent biofilm was fixed with 100 μL of 99% methanol for 15 minutes, washed with DI water, and air-dried at room temperature. Biofilms were stained with 100 μL of 0.2% crystal violet for 15 minutes, washed with running water, and dried at room temperature for 30 minutes. Once dry, wells were solubilized with 150 μL of 15% acetic acid to release the crystal violet staining into solution. Quantification was performed by measuring absorbance at 570 nm using a microplate reader.

### Confocal imaging

Briefly, cultures were prepared as outlined above with 1 mL of bacterial culture inoculated into each chambered glass slide. Biofilms were stained with BacLight™ Red Bacterial Stain (Thermo Fisher Scientific) using a 200 μM working solution, with 2 μL added per 1 mL of GBS sample. Samples were incubated with the stain in the dark at 37°C overnight. Following incubation, the staining solution was carefully removed and then gently washed with 500 μL PBS to remove excess stain before imaging. Biofilms were imaged using a Leica Stellaris confocal laser scanning microscope (Leica Microsystems). Z-stack images of the biofilm were acquired using identical imaging parameters for all samples to compare between non-desiccated A909 and D30 strains. Images were processed using LAS X software (Leica Microsystems). Mean fluorescence intensity was measured using ImageJ software (version 1.54g).

### Adherence and invasion assay

Adherence and invasion assays were performed using human vaginal epithelial cells (VK2, ATCC). VK2 cells were seeded in 24-well tissue culture plates at approximately 2.5 x 10^5^ per well and grown to 90-100% confluence in 24-well tissue culture plates. Frozen stocks of the non-desiccated and desiccated A909 strains were grown to mid-log phase (OD_600_ of 0.4), resuspended in fresh keratinocyte serum free cell culture medium (KSFM). Bacterial suspensions were diluted 1:10 into KSFM and 50 μL was added to each well containing 450 μL KSFM, approximately 5 x 10^5^ CFU/well, corresponding to an MOI of about 2:1. Plates were centrifuged at 100 × g for 5 minutes and incubated at 37°C with 5% CO_2_. For the adherence assay, infected monolayers were incubated for 30 minutes and washed five times with 0.5 mL PBS to remove non-adherent bacteria. Cells were released from the plate with 100 μL trypsin, lysed with 400 μL 0.025% Triton X-100, serially diluted, and plated on THB agar for CFU/mL enumeration. For invasion assays, infected monolayers were incubated for 2 hours at 37°C with 5% CO_2_, washed, and treated with 5 μg/mL of penicillin and 100 μg/mL of gentamicin containing medium for 1 hour to kill extracellular bacteria. Cells were then washed three times, trypsinized, lysed with 0.025% Triton X-100, serially diluted, and plated for CFU enumeration. Results are expressed as the percentage of the input inoculum recovered from adherence or invasion.

### Stress survival assay

Frozen stocks of the non-desiccated and desiccated A909 strains were grown to mid-log and normalized in PBS. For UV exposure, the bacterial cells were exposed to UV for 15 minutes, then diluted and plated on THB agar to determine viability. For growth-based stress assays, 200 μL of 1 x 10^6^ CFU/mL GBS were added to each well of a 96-well plate in the following conditions: I) 3% ethanol stress (Bio Lab Ltd) and II) 100 ppm sodium hypochlorite stress prepared from a 5% stock solution (LabChem). Samples were plated for viability at time 0 to prior to treatment and were plated at 6 hours and 18 hours after treatment, respectively. All experiments were performed in at least three independent biological replicates.

### Murine model of vaginal colonization

Vaginal colonization studies were performed using a previously established murine model [4, 23]. Female 8-week-old ICR mice (Envigo) were synchronized in estrus cycle with 0.5 mg of β-estradiol in 100 μL sesame oil administrated via intraperitoneal a day prior to inoculation. On experimental day 0, animals were inoculated intravaginally with approximately 1 x 10^7^ CFU/10 μL with either the non-desiccated A909 or D30 strain. To assess bacterial viability in the vaginal lumen over time, vaginal swabs were collected daily, and samples were serially diluted and plated on StrepB CHROMagar (Fisher Scientific), to select for GBS. On day 5 post-inoculation, mice were humanely euthanized, and the vagina, cervix, and uterus were harvested. Tissues were homogenized, serially diluted, and plated on CHROMagar to determine GBS burden. CFUs were normalized to tissue weight.

### Murine model of hematogenous systemic infection

For the systemic model of infection, we used the previously established murine model [24]. Female 8-week-old female ICR mice (Envigo) were used for this experiment. Animals were anesthetized with isoflurane and the eye was covered with a drop of proparacaine solution and inoculated via retro-orbital injection with approximately 2-3 x 10^8^ CFU in a 100 μL volume in PBS of either A909 or D30 and monitored for 83 hours. Disease progression of each animal was evaluated based on scoring criteria that is approved under UTK IACUC protocol no. 2937-1222 that assesses gait, hunching, signs of pain, grooming, scruffiness of fur, breathing patterns, and onset of neurological symptoms. Each criterion is measured on a 0-3 scale and animals were monitored continuously after the onset of symptoms. Upon reaching a moribund state or a health score of 12. Animals were humanely euthanized and blood, heart, brain, and lung tissue were harvested. Tissues were homogenized and plated on THA to assess bacterial burden. CFUs were normalized to tissue weight.

### Whole genome sequencing and variant analysis

Stocks of the non-desiccated and desiccated A909 strains were plated and 5 replicate colonies were sub-cultured to OD_600nm_ of 0.4 and DNA was extracted using Quick DNA Miniprep Plus Kit (Zymo research) per manufacturer’s instructions and quantified by Nanodrop. Samples were diluted and submitted to SeqCoast Genomics for Illumina sequencing. Illumina paired end sequencing was performed, generating approximately 200 Mbp and 1.3 million reads per sample. Sequence reads were trimmed using Trimmomatic v0.39 [25] and mapped to the *S. agalactiae* A909 reference genome (GCF_000012705.1) using Bowtie2 v2.5.3 [26]. Genetic variants were identified using breseq v0.38.3 through the SeqCoast prokaryotic variant-calling pipeline in population mode identifying mutations detected in the desiccated population to the non-desiccated population [27].

### Statistical analysis

Data analysis was performed using GraphPad Prism version 9.5.1 for iMac, GraphPad Software, San Diego, California, USA. Statistical significance and specific tests are indicated in the figure legends.

## Discussion

Although GBS is a leading cause of neonatal sepsis and meningitis, relatively little is known about its survival and persistence outside the host [2, 4]. Environmental persistence has been increasingly recognized as an important contributor to transmission of several pathogens, allowing bacteria to survive on abiotic surfaces for prolonged periods before infecting a susceptible host [28-30]. Closely related streptococci, *Streptococcus pneumoniae* and Group A *Streptococcus* (GAS), have been shown to similarly survive several weeks of desiccation and on contaminated environmental surfaces before transmission [21, 31]. Studies have shown that invasive GBS infection in one infant born from multiple births is readily transmitted to infect the other(s) infant siblings supporting contact-mediated transmission, and studies have reported that GBS from hospital environments, medical equipment, and environmental contamination contributes to transmission in neonatal healthcare settings [14-16, 32-34]. Our findings demonstrate that GBS A909 remains viable in PBS without the addition of nutrients for up to 30 days at room temperature and remains viable for more than 80 days under desiccated conditions. One notable phenotypic change observed following GBS desiccation was enhanced biofilm formation, which has been demonstrated as an important adaptation to promote survival during environmental stress [35]. Previous work in *S. pyogenes* and *S. pneumoniae* shows that biofilm formation [14-16] promotes survival under desiccating conditions [36]. Our data are complementary and show that prior stress by desiccation results in an increase in GBS biofilm formation, linking the two clinically relevant phenotypes across diverse streptococcal pathogens [21, 36]. Importantly, we observed survival during desiccation and enhanced biofilm formation in multiple clinical GBS isolates and across capsular serotypes Ia and III, suggesting the significance of this adaptive response is not restricted to a single strain or serotype. This persistence was not accompanied by increased tolerance to the disinfectant and environmental stresses tested, though we tested for tolerance in planktonic grown GBS. It would be interesting to test these similar treatment conditions in biofilm-grown GBS to determine if the increased biofilm observed in our desiccated strains impacts resistance to disinfectants. This information would be significant for ensuring that infection-control practices remain effective for limiting environmental GBS and reducing opportunities for transmission.

Whole genome sequencing of our desiccated GBS population identified enriched mutations in genes encoding proteins associated with cell surface and envelope homeostasis that promote GBS survival during outgrowth following desiccation and were absent in the non-desiccated A909 strain. Particularly notable were multiple insertion–deletion events within the coding region of the capsular polysaccharide synthesis gene, *cpsH*. Capsular architecture is closely linked to surface-associated adhesion in GBS and plays an important role in virulence and host interactions [8, 37, 38]. Alterations in capsule production can also influence biofilm development; experimentally increasing capsular polysaccharide production in *S. agalactiae* reduced biofilm formation, adhesion, and invasion, demonstrating that changes in capsule abundance can substantially alter surface-associated phenotypes and the corresponding capsule-associated proteins [39]. The potential relevance of capsule-associated pathways to desiccation is also supported by studies in *Staphylococcus aureus*, where desiccation significantly enhanced expression of capsular polysaccharide synthesis genes and disruption of capsule production reduced survival during drying [40].

Additional mutations were detected in *fbsA*, which encodes a cell-wall-anchored fibrinogen-binding protein that promotes GBS adherence to epithelial cells. Variation in *fbsA* has been associated with differences in fibrinogen-binding capacity among clinical GBS isolates [41, 42]. Mutations were also detected in *liaF, pbp3*, and D-Ala-D-Ala ligase (*ddl*), collectively pointing toward pathways involved in cell-envelope stress responses and peptidoglycan homeostasis. LiaF is part of the *liaFSR* cell-envelope stress response system, which contributes to maintenance of GBS cell-wall integrity [43], whereas PBP3 and Ddl participate in peptidoglycan synthesis through distinct steps in cell-wall assembly [44-46]. Notably, pbp3 (A123T) and ddl (E262K) are non-synonymous substitutions that alter the encoded amino acid sequence. Maintenance of cell-wall integrity has emerged as an important determinant of desiccation tolerance in other Gram-positive bacteria [21, 31, 47]. In *S. pneumoniae*, disruption of PBPs involved in cell-wall maturation reduced survival during desiccation, demonstrating that PBP-mediated cell envelop homeostasis can influence desiccation tolerance [48]. Studies in other bacteria have shown that loss of type A PBPs has been characterized as leading to decreased cell wall stiffness and fewer peptidoglycan cross-links [49, 50]. Moreover, cell-wall synthesis influences GBS biofilm development, as exposure to the PBP-targeting antibiotic penicillin reduces biofilm biomass and thickness [46] while disruption of D-alanine metabolism in *S. mutans* results in cell-wall defects accompanied by reduced biofilm formation and extracellular polysaccharide production [44, 45, 51, 52]. Overall, these findings suggest that alterations in peptidoglycan homeostasis may contribute to the enhanced biofilm formation and environmental persistence observed following GBS desiccation.

Additional mutations in the desiccated population were detected in genes involved in carbohydrate and energy metabolism, and nutrient transport. These included variants in a phosphotransferase system, *glgA, nikA*, glucosamine-6-phosphate deaminase, among others. Although the functional significance of these mutations remains unknown, they collectively point toward metabolic adaptation as a potential component of long-term environmental persistence. This interpretation is consistent with our observation that GBS survives not only prolonged desiccation but also extended nutrient deprivation. In *S. pneumoniae*, Tn-seq identified multiple carbohydrate transporters, carbohydrate catabolic genes, and a putative metal transporter whose disruption altered desiccation fitness, while transposon screening in *L. monocytogenes* similarly implicated energy metabolism and membrane transport in survival during drying [48, 53]. These observations suggest that successful persistence of GBS under environmental stress likely depends on coordinated adaptation of both metabolic and cell-envelope functions. Further studies are needed to determine whether these mutations contribute directly to desiccation survival and the associated phenotypic changes.

Environmental persistence may be particularly relevant to LOD in neonatal healthcare settings, for which horizontal transmission has become increasingly recognized [54-56]. Previous work identified four distinct clusters of invasive GBS LOD cases within a single NICU over two years, and whole-genome sequencing revealed genetic links between 11 of 12 invasive isolates to at least one other isolate in the same clinical setting [57]. Other studies have reported transmission from healthcare or community sources, including GBS nosocomial acquisition or mother-to-baby transmission after delivery, with significantly higher rates of neonatal colonization specifically of the LOD-associated hypervirulent CC17 clade detected further from the time of birth [58-60]. The significance of environmental persistence depends ultimately on whether bacteria surviving outside the host maintain infectivity [47]. Here we demonstrate that desiccated GBS retain virulence compared to non-desiccated control GBS strains, with desiccated and non-desiccated GBS colonizing the vaginal lumen to comparable levels, though we detected enhanced reproductive tissue invasion and enhanced systemic pathogenesis in desiccated GBS compared to the non-desiccated control strain. Together, these findings indicate that GBS persistence during nutrient deprivation and desiccation does not result in a loss of infectivity and instead may prime GBS for subsequent host infection or select for a GBS population that is more resistant to clearance by the host immune response. This may be particularly relevant in neonatal and other healthcare settings, where opportunities for indirect transmission pose serious threat to vulnerable patients.

## Acknowledgements

We thank SeqCoast Genomics (Porsmouth, NH, USA) for sequencing and bioinformatics support. We also thank Drs. Andrew Monteith and Todd Reynolds for their insight on this project.

## Figure Legends

**Supplementary Figure 1. Impacts of desiccation on GBS virulence**

**A)** Comparison of biofilm formation of control and desiccated GBS strains from representative serotypes. Statistical analysis was performed using two-way ANOVA with Sídák’s multiple comparisons test. **B)** Representative images of hemolytic potential of non-desiccated A909 and A909 D30 on 5% sheep’s blood agar. **C)** Adherence to VK2 vaginal epithelial cells. Survival of non-desiccated A909 and D30 following general stresses including **D)** UV exposure, **E)** ethanol stress, or **F)** sodium hypochlorite exposure. **G)** Invasion of VK2 vaginal epithelial cells. Data are presented as mean ± SEM from at least three independent biological replicates. Statistical analyses for panels C and G were performed using an unpaired t-test with *p* values indicated on individual figures.

