## Supplemental Figure 1 for "Desiccation stress enhances virulence potential of group B *Streptococcus*"

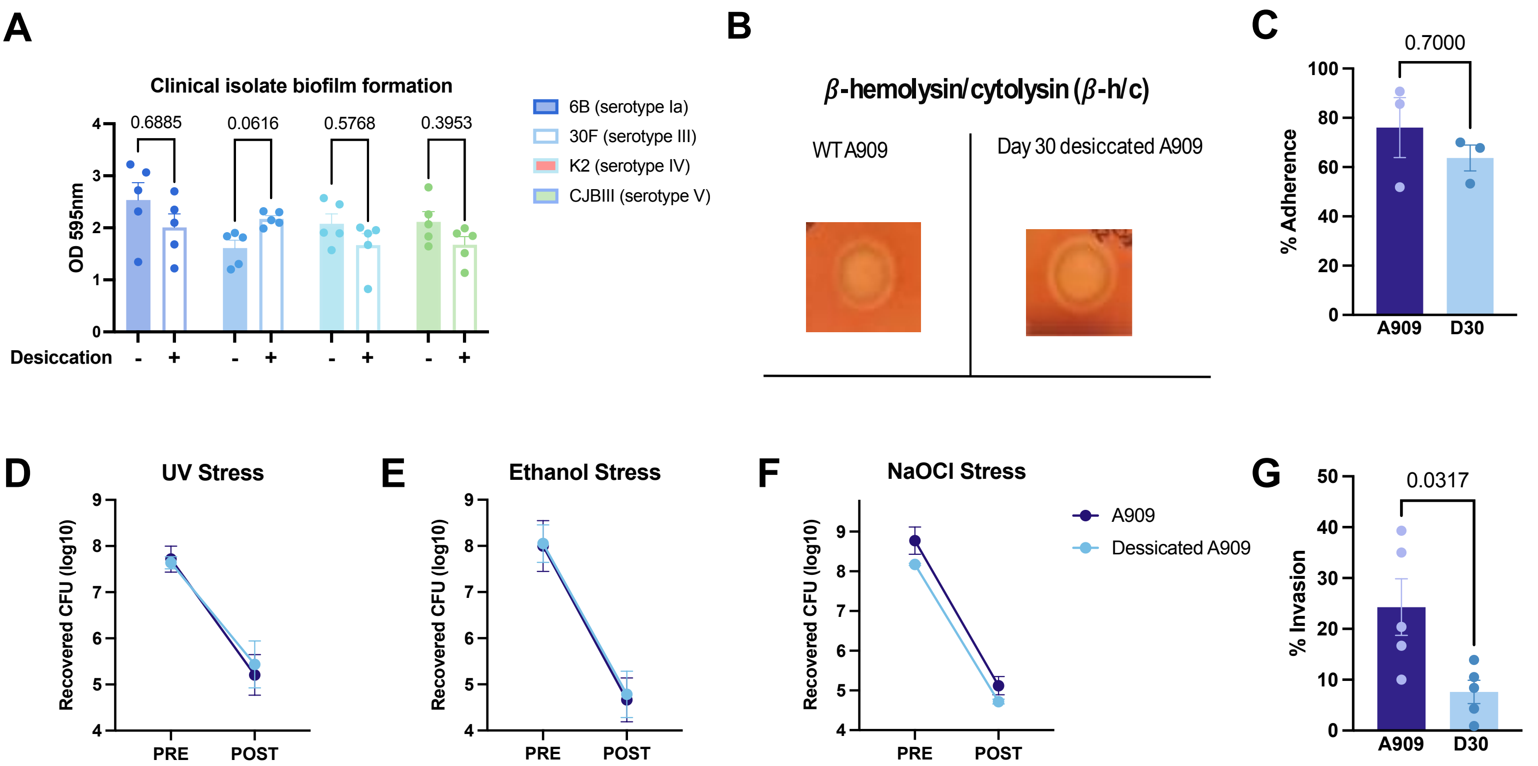
A
Clinical isolate biofilm formation
OD 595nm
0.6885
0.0616
0.5768
0.3953
Desiccation
6B (serotype Ia)
30F (serotype III)
K2 (serotype IV)
CJBIII (serotype V)
B
β-hemolysin/cytolysin (β-h/c)
WT A909
Day 30 desiccated A909
C
0.7000
% Adherence
A909
D30
D
UV Stress
Recovered CFU (log10)
PRE
POST
E
Ethanol Stress
Recovered CFU (log10)
PRE
POST
F
NaOCl Stress
Recovered CFU (log10)
PRE
POST
A909
Dessicated A909
G
0.0317
% Invasion
A909
D30
